# Differential impact of maternal overnutrition and undernutrition on offspring glucose homeostasis

**DOI:** 10.1101/2020.12.03.410720

**Authors:** Praise B. Adekunbi, Abimbola O. Ogunsola, Daniel A. Adekunbi

## Abstract

To test the hypothesis that maternal undernutrition exerts a greater effect on offspring metabolic function compared to maternal overnutrition, female rats (n=10 per group) were subjected to a high calorie diet or 50 % global nutrient restriction relative to control rats on standard rat chow, for 8 weeks prior to pregnancy and during pregnancy. Birth weight was determined on the day dams were found with pups. At 3 months of age, offspring’s fasting blood glucose, serum insulin and triglyceride levels as well as glucose tolerance and insulin sensitivity were determined. A 50 % calorie restriction caused a significant weight loss in the under-nourished dams but those on high calorie diet had similar body weight as control rats. Maternal overnutrition and undernutrition significantly lowered birth weight, indicating intra-uterine growth restriction in these animals. Fasting blood glucose was significantly higher in female offspring of over-nourished dams, but neither maternal overnutrition nor undernutrition affected offspring’s glucose tolerance. Male offspring of dams exposed to maternal overnutrition or undernutrition had a significantly higher insulin level compared to control, whereas female offspring were unaffected. The development of hyperinsulinaemia in male offspring of undernourished dams was accompanied by reduced insulin sensitivity. This study demonstrates that early-life exposure to two extreme ends of the nutritional plane is associated with similar birth weight outcome but different metabolic phenotype in adulthood. Evidence of insulin resistance only in male offspring of under-nourished dams indicates differences in sex-specific metabolic effect of maternal undernutrition compared to overnutrition.

## 1.0. Introduction

Developmental programming relates to the process whereby an environmental factor including altered nutrition during the developmental period cause alterations in tissue structure and function leading to disease susceptibility in adult life (Segovia *et al.*, 2014). Studies dating back to more than three decades have shown an association between low birth weight arising from maternal under-nutrition and incidences of hypertension, stroke and metabolic syndrome in adulthood (Barker and Osmond, 1986; Barker *et al.*, 1989), which has led to the term developmental origin of health and diseases (DOHaD) (Hanson and Gluckman, 2014). The concept of DOHaD also operates in the context of excessive and rich nutritional environment (Segovia *et al.*, 2014), given that maternal obesity predisposes the offspring to develop obesity, type 2 diabetes and cardiovascular diseases (Catalano *et al.*, 2009; Labayen *et al.*, 2010). A maladaptive intra-uterine environment permanently alters the structure of vital organs of a developing foetus to ensure immediate survival of the suboptimal environment (Tarry-Adkins and Ozanne, 2017). These vital organs include the heart, liver, kidney, pancreas as well as adipose tissue and alterations in their growth, vasculature or metabolic activities underpin disease risk in adulthood.

Undernutrition and overnutrition during pregnancy remain a public health concern in both developing and developed economies (Deputy *et al.*, 2018; Dadi and Desyibelew, 2019; Koletzko *et al.*, 2019). Clearly, there is a link between the two extreme nutritional statuses during development on later life health of the offspring (Odhiambo *et al.*, 2020). This study seeks to compare metabolic outcomes associated with maternal overnutrition and undernutrition in adult offspring and determine the pattern of sexual dimorphic response arising from such early life nutritional exposure. To test the hypothesis that maternal undernutrition exerts a more debilitating programming effect on offspring metabolic function over maternal overnutrition, we determined birth weight outcome as a measure of foetal development as well as glucose homeostasis, insulin and triglyceride concentrations in adult offspring.

## 2.0. Methods

### 2.1. Animals

Thirty (30) female and ten (10) male Sprague-Dawley rats averagely weighing 125 g were used in this study. The female rats were randomly divided into three groups, namely control and overnutrition and undernutrition. The animals were kept in the animal facility of Babcock University under controlled environmental conditions (12 h light, 12 h dark cycle; temperature 24 ^o^C ± 2 ^o^C). All guidelines with the use and care of laboratory animals were strictly adhered to and in accordance with the criteria outlined by the National Academy of Science published by the National Institute of Health (NIH, 1985).

### 2.2. Experimental design

Control rats received a standard rat chow which comprised 7% simple sugars, 3 % fat, 50 % polysaccharide, 15 % protein, energy, 3.5 kcal/g. The animal exposed to overnutrition received an energy-dense high fat diet (20 % animal lard, 10 % simple sugar, 23 % protein, energy, 4.5 kcal/g, Special Dietary Services, Essex UK) and sweetened condensed milk (Nestlé, Vevey, Switzerland) diluted to equal proportion (1:1) in drinking water. The undernutrition group received 50 % of *ad libitum* food consumed by control group. The female rats were on the assigned dietary regime for 8 weeks before mating and throughout pregnancy, with body weight monitored weekly. Proven fertile males on control diet were used for mating.

On the day dams were found with pups, birth weight of each pup was determined using a sensitive weighing scale. The litter size was standardized two days after birth to 6 pups (3 males, 3 females) per dam across the experimental groups to ensure equal nutritional access during suckling. Dams with less than 6 pups or odd sex ratio were not included in the study. All offspring were weaned and grouped into separate cages according to their gender on postnatal day (pnd) 21 and maintained on a standard rat chow until postnatal day (pnd) 90 when glucose tolerance and insulin sensitivity tests as well as insulin and triglyceride concentrations were determined. To avoid litter bias on study outcome, only two males and 2 females from each litter were used at a given time for the various endpoints examined.

### 2.3. Blood glucose and glucose tolerance test

Male and female offspring of control, over-nourished and under-nourished dams were fasted overnight before blood samples were collected from the tail vein for glucose measurement using a glucose monitoring system (Accu-check glucometer, Roche, Germany). Oral glucose tolerance test was carried out to determine the regulation of glucose metabolism in these animals. Timecourse changes in blood glucose level was determined with a glucose load of 2 g/kg body weight. Blood glucose concentrations were determined from the tail vein at 30, 60, 90, and 120 min post glucose gavage. The area under the glucose curve (AUC) was also calculated.

### 2.4. Insulin tolerance test

To determine insulin sensitivity in rat exposed to maternal overnutrition and undernutrition, rats that were fasted for 6 h received intraperitoneal injection of human insulin solution (Humulin, 0.75U/kg body weight). The blood glucose concentration was monitored before (0 min) and 15, 30, 60, 90, and 120 min after insulin injection. The AUC for the time-course blood glucose during the insulin tolerance test was calculated.

### 2.5. Serum insulin and triglyceride levels

A subset of male and female offspring of control, over-nourished and under-nourished dams were culled by cervical dislocation on pnd 90 (n=6 per sex) and blood was collected via cardiac puncture. The blood samples were centrifuged at 3000 revolutions per minute (rpm) for 10 minutes to obtain serum for estimation of insulin and triglyceride levels. Serum insulin level was determined using the enzyme linked immunosorbent kit (RayBio^®^Rat Insulin) obtained from RayBiotech Inc (Norcross, USA). The sensitivity of the assay was 5 μIU/ml. The intra-assay and inter-assay variation were 10 % and 12 % respectively. Serum triglyceride level was estimated using kits purchased from Randox Laboratories Limited (Crumlin, United Kingdom) with measurement carried out with an automatic biochemistry analyzer (BT, 2000 Plus, Germany).

### 2.6. Statistical analysis

All results are presented as the mean ± standard error of mean (SEM). Data analyses were carried out using GraphPad Prism Software (GraphPad, Inc., La Jolla, CA, USA). Analysis of birth weight data was performed using one-way analysis of variance (ANOVA) with post hoc Tukey’s multiple comparison test. Two-way ANOVA was used to determine the interaction between offspring nutritional exposure and sex on body weight and metabolic parameters examined. significance was set at p < 0.05.

## 3.0. Results

### 3.1. Maternal body weight before and during pregnancy following dietary regimen

Exposure of female rats to a high calorie diet elicited changes in body weight that was similar to those on control diet. However, a 50 % calorie restriction for 8 weeks prior to and throughout pregnancy caused a significant decrease in body weight (Table 1).

**Table 1:** Body weight before and during pregnancy in over-nourished and under-nourished dams

|  | Timeline | Control | Overnutrition | Undernutrition |
| --- | --- | --- | --- | --- |
| **Body weight before pregnancy (g)** | Baseline | 127.2 ± 3.50 | 126 ± 4.38 | 135.6 ± 6.09 |
|  | Week 8 | 165.2 ± 4.99 | 181.7 ± 9.51 | 129.0 ± 7.79 *# |
| **Body weight during pregnancy (g)** | Week 1 | 177.2 ± 6.28 | 186.2 ± 10.38 | 128.7 ± 5.48 *# |
|  | Week 2 | 209.4 ± 7.91 | 202.5 ± 11.89 | 121.2 ± 9.08 *# |
|  | Week 3 | 243.8 ± 7.39 | 220.3 ± 14.59 | 151.3 ± 11.57 *# |

Data are expressed as mean±SEM, n=6, \*p<0.05 compared with control, #p<0.05 compared with overnutrition.

### 3.2. Effects of maternal overnutrition and undernutrition on birth weight

The birth weight of offspring of control, over-nourished and under-nourished dams shown in Figure 1 were pooled data from male and female pups. The birth weight of pups of dams exposed to overnutrition or undernutrition was significantly lower than birth weight of control dams, which indicates the dietary exposure caused intrauterine growth restriction (IUGR). The severity of the IUGR was higher in undernutrition group given their significantly lower birth weight compared with overnutrition group.

**Fig. 1:**
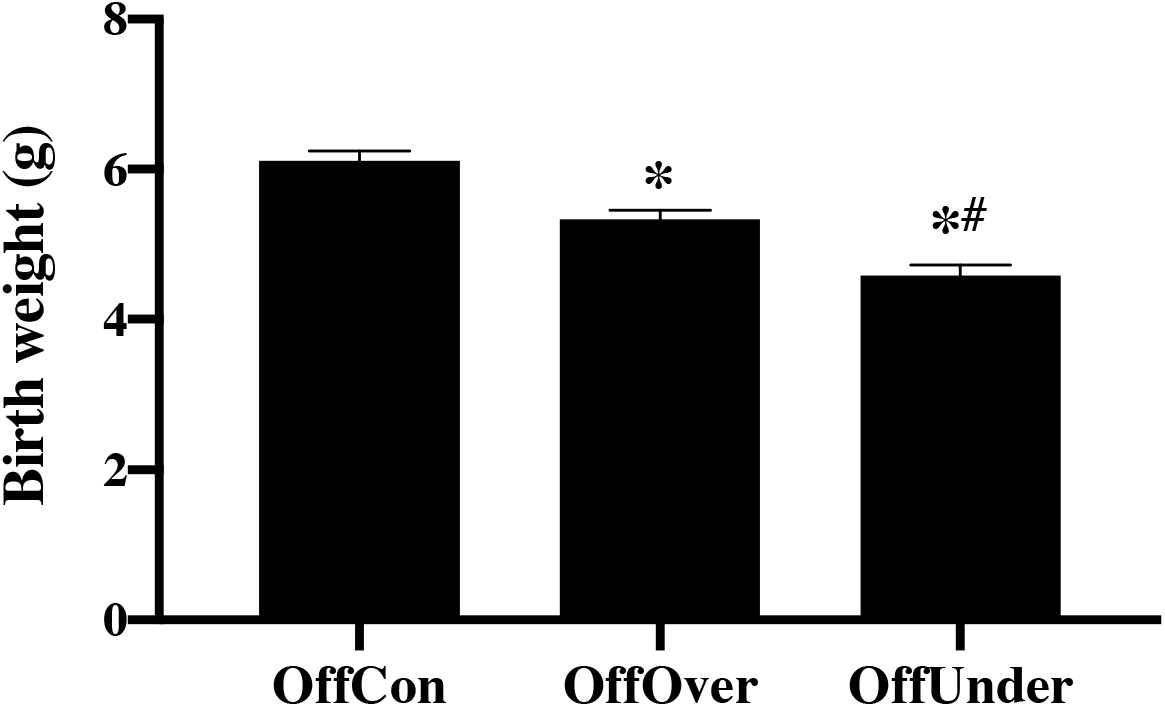
Birthweight in offspring of control, over-nourished and under-nourished dams. Results are presented as mean ± SEM, n=27-35. *p <0.05 compared with OffCon, #p <0.05 compared with OffOver. Abbreviation: OffCon - offspring of control dams; OffOver - offspring of overnourished dams; Offspring of under-nourished dams.

### 3.3. Effects of maternal overnutrition and undernutrition on post-weaning body weight

Offspring postnatal weight was determined at 3 months of age (Postnatal day 90), The differences in weight at birth did not persist into adulthood indicating a catch up in growth rate in rats subjected to *in utero* overnutrition or undemutrition. Meanwhiie, male offspring had significantly higher body weight compared with female (Figure 2) corroborating the well-known adult body weight differences in male and female rats.

**Fig. 2:**
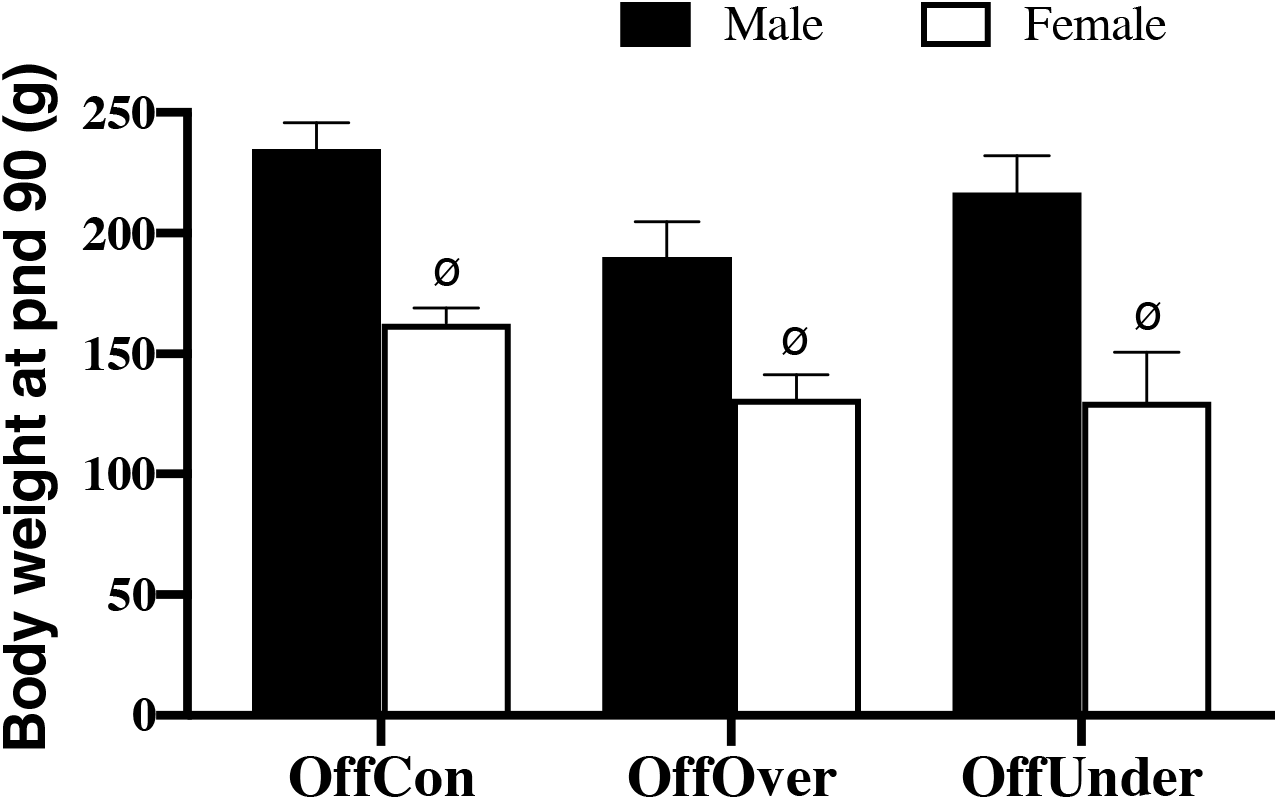
Body weight of offspring of control, over-nourished and under-nourished dams on postnatal day 90. Results are expressed mean ± SEM, n=6. ^ø^p <0.05 compared with female OffCon. Abbreviation: OffCon - offspring of control dams; OffOver - offspring of overnourished dams; Offspring of under-nourished dams.

### 3.4. Effects of maternal overnutrition and undernutrition on offspring fasting blood glucose level

As shown in figure 3, there were no significant alterations in mean fasting blood glucose (FBG) level of adult male offspring of over-nourished and under-nourished dams (OffOver and OffUnder respectively) when compared with offspring of control dams (OffCon). FBG tended to be higher in male OffOver compared with OffCon. In the female, fasting blood glucose level was significantly higher in OffOver than OffCon and OffUnder.

**Fig. 3:**
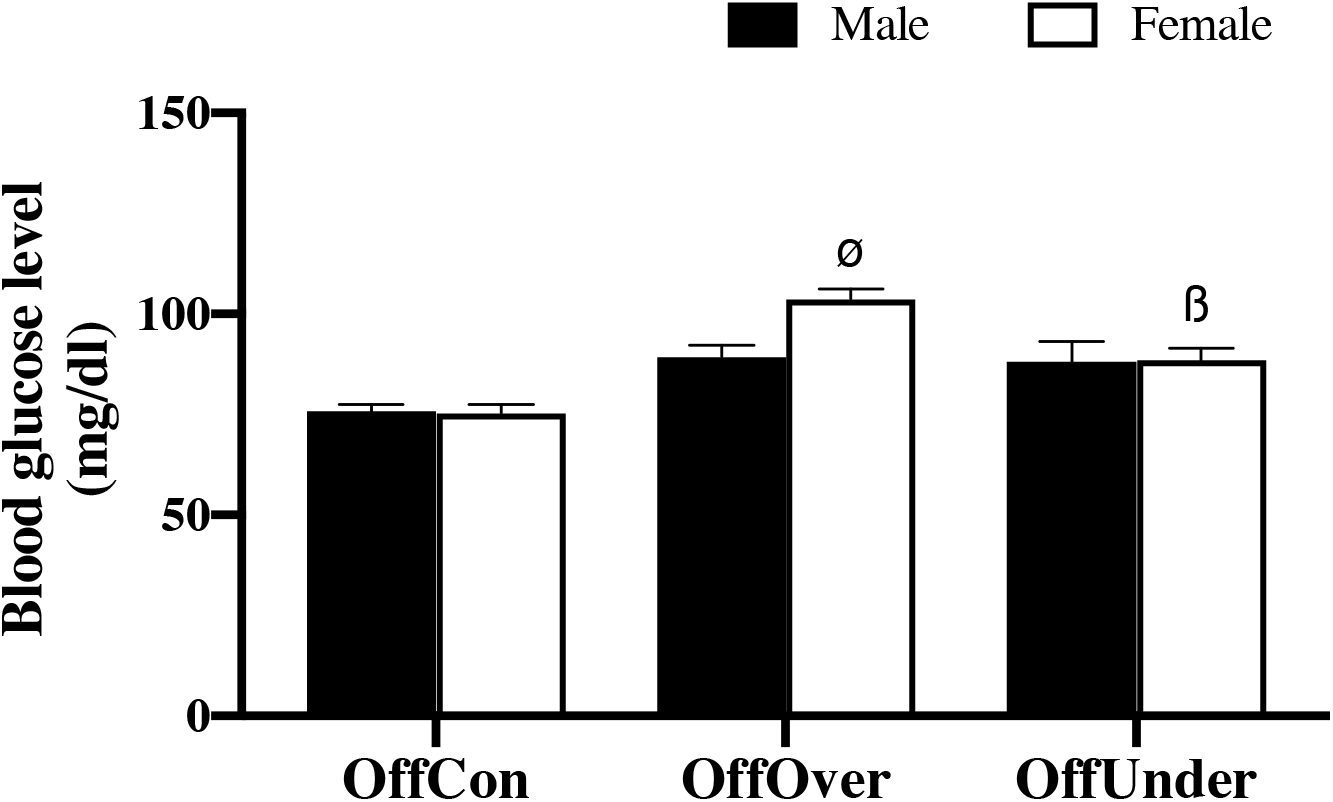
Fasting blood glucose levels in offspring of rats exposed to maternal overnutrition and undernutrition on postnatal day 90. Data expressed mean ± SEM, n=6. ^ø^p <0.05 compared with female OffCon, ^ß^p <0.05 compared with female OffOver. Abbreviation: OffCon - offspring of control dams; OffOver - offspring of over-nourished dams; Offspring of under-nourished dams.

### 3.5. Effect of maternal overnutrition and undernutrition on offspring glucose tolerance

To investigate the programming effect of maternal overnutrition and undernutrition on glucose tolerance, the offspring were challenged with a 2 g/kg glucose load. The time-course response to the glucose challenge in male OffOver and OffUnder was similar to that of male OffCon leading to a comparable AUC during the oral glucose tolerance test. In the female, at 30 min post-glucose challenge, the blood glucose level of OffCon was significantly higher than OffOver and OffUnder. At subsequent time points, the blood glucose level was similar among the experimental groups except at 180 min where the blood glucose level of OffCon was lower than that of OffOver. The area under the glucose curve was not significantly different for all comparisons made in female offspring (Figure 4).

**Fig. 4:**
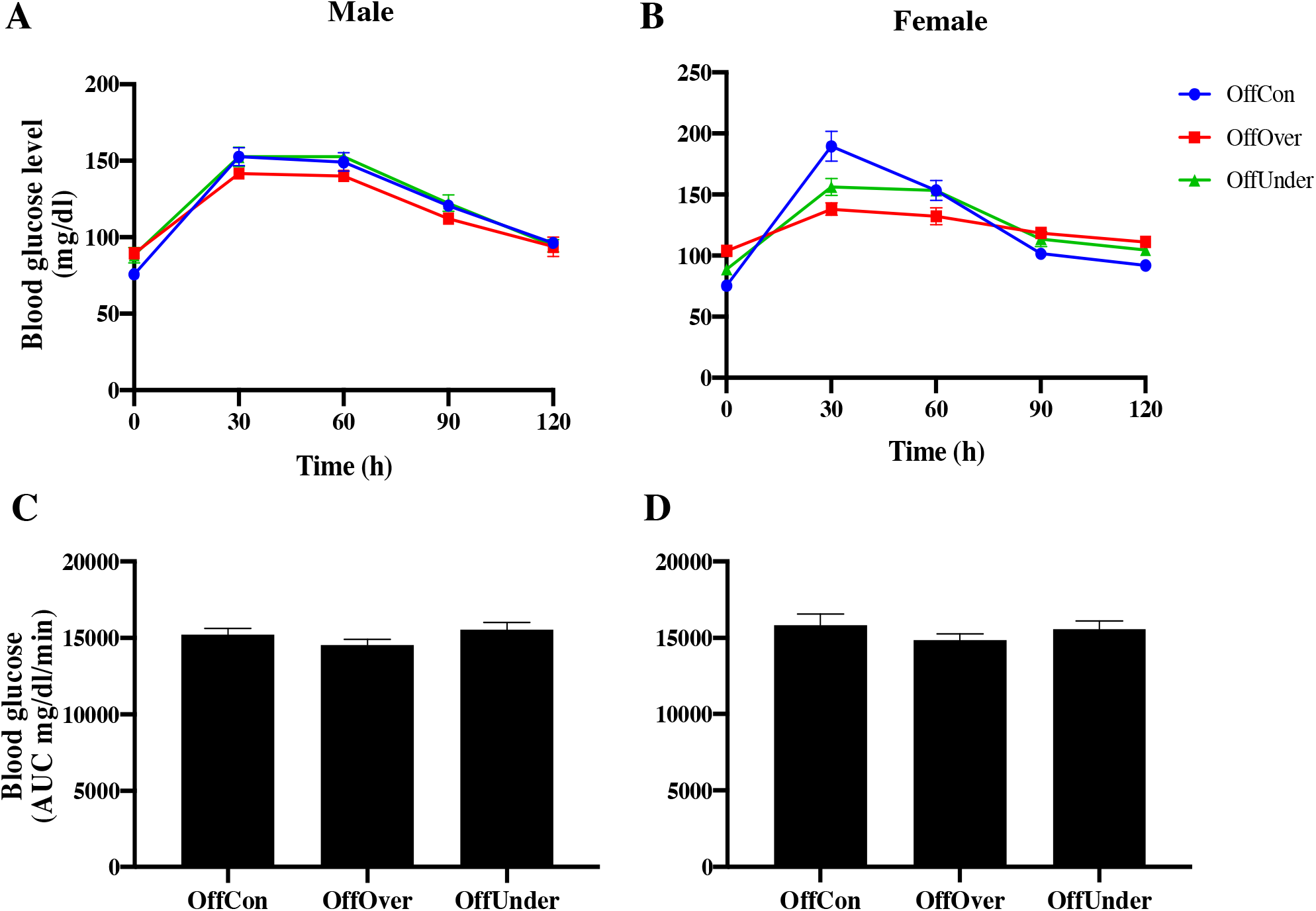
Glucose response curves in male (A) and female (B) offspring of rats exposed to maternal over-nutrition and under-nutrition during oral glucose tolerance test with area under the curve (AUC) in male (C) and female (D) offspring. Data expressed mean ± SEM, n=6. Abbreviation: OffCon - offspring of control dams; OffOver - offspring of over-nourished dams; Offspring of under-nourished dams.

**Fig. 5:**
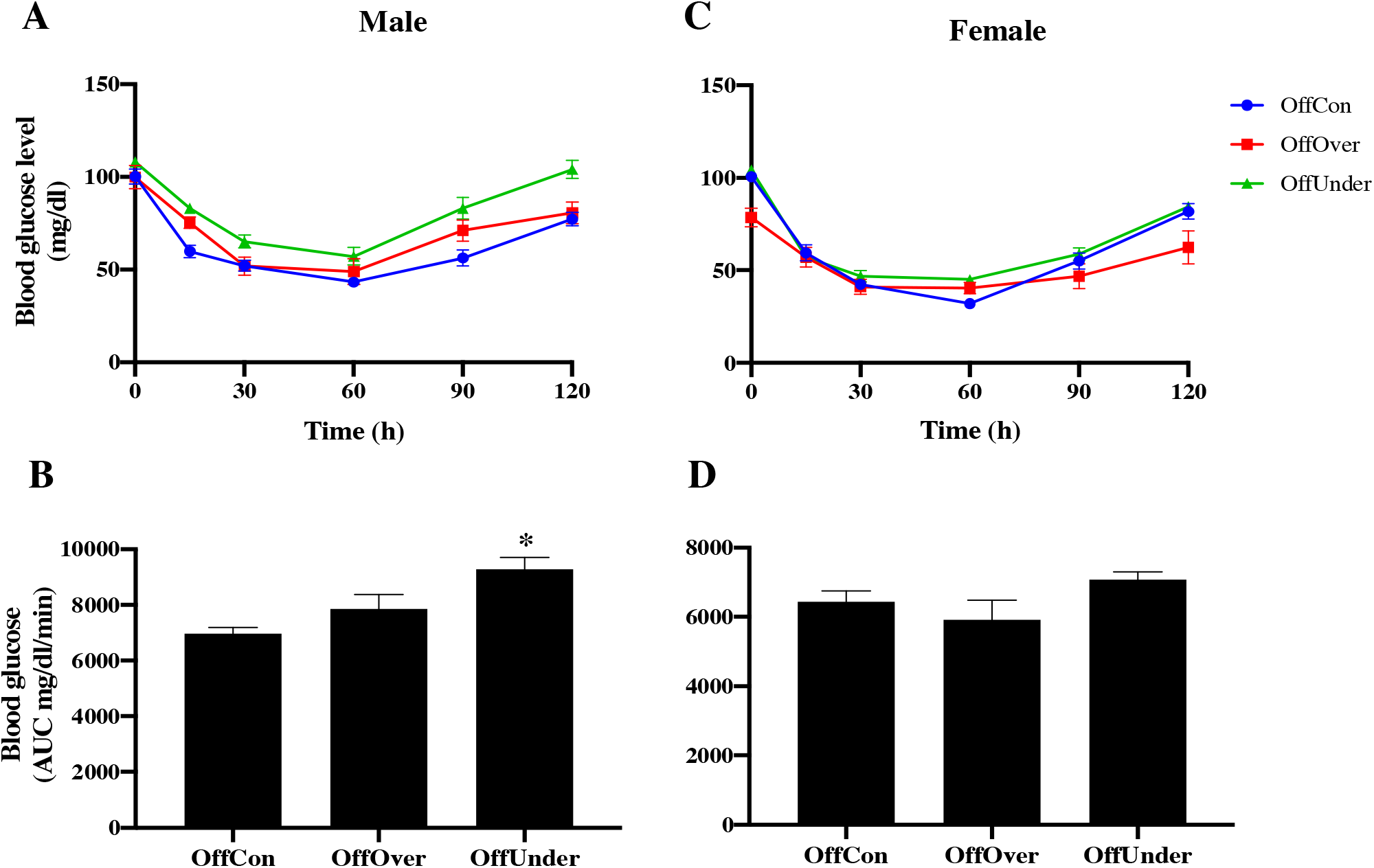
Glucose response curves during insulin tolerance test in male (A) and female (B) offspring of rats exposed to maternal overnutrition and undernutrition. Area under the curve (AUC) derived from glucose response to insulin challenge in male (C) and female (D) offspring. Data expressed mean ± SEM, n=6. *p <0.05 compared with male OffCon. Abbreviation: OffCon - offspring of control dams; OffOver - offspring of over-nourished dams; Offspring of undernourished dams.

**Fig. 6:**
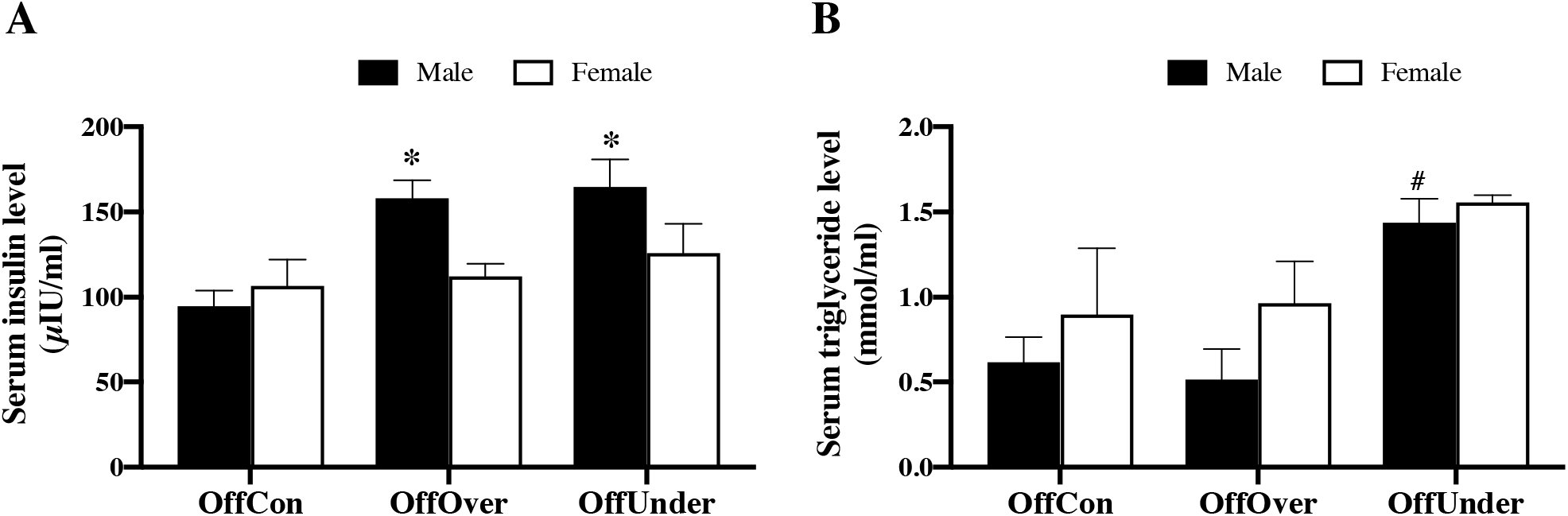
Serum insulin (A) and triglyceride (B) levels in male and female offspring of rats exposed to maternal overnutrition and undernutrition. Results are presented as mean ± SEM, n=27-35. *p <0.05 compared with male OffCon, #p <0.05 compared with male OffOver. Abbreviation: OffCon - offspring of control dams; OffOver - offspring of over-nourished dams; Offspring of under-nourished dams.

### 3.5. Effect of maternal overnutrition and undernutrition on insulin sensitivity in the offspring

The time-course blood glucose level during the insulin tolerance test (ITT) as well blood glucose AUC during the ITT in male and female offspring of rat exposed to over-nutrition or undernutrition is presented in figure 2. As expected, following intraperitoneal insulin injection, there was a drastic fall in blood glucose level as early as 15 min post-injection. Meanwhile, after 2h insulin injection, blood glucose level was significantly higher in male offspring of undernourished dams compared to both control and overnutrition group indicating reduced sensitivity to insulin challenge. The AUC during the ITT was significantly higher in male OffUnder compared to OffCon and OffOver. In the female, the blood glucose response during the insulin tolerance test as well as the AUC were similar regardless of early-life nutritional status.

### 3.6. Effects of maternal overnutrition and undernutrition on offspring insulin level

In male offspring of rats exposed to maternal overnutrition or undernutrition, serum insulin level was significantly higher than control offspring but not in females (Figure 3A). Serum triglyceride level was significant higher in male OffUnder when compared with male OffOver. In the female, the level of triglyceride was similar among the experimental groups (Figure 3B)

## 4.0. Discussion

Foetal growth limitation due to a poor intra-uterine environment is related to the development of metabolic diseases in adult life (Chaudhari *et al.*, 2017). In the context of poor nutritional environment, it is possible that the available nutrients are redirected towards the development of critical organs such as the brain and heart at the expense of less critical ones like the pancreas and kidney, leading to development of metabolic diseases during postnatal life (Devaskar *et al.*, 2016).

The present study demonstrates that two disparate nutritional environment elicits similar adverse foetal growth condition resulting in low birth weight. We observed that the impact of maternal nutrition on birth weight seems to be more severe in the undernourished group compared to overnutrition, which may be due to availability and utilization of nutrients. Nevertheless, there is a possibility of a shared mechanistic pathway in the placenta to cause foetal growth restriction in an excessive or deficient nutritional condition. While the reduced birth weight in offspring of under-nourished dams corroborate previous studies (Ramírez-López *et al.*, 2017), there are conflicting reports in the literature on the impact of maternal overnutrition on offspring birth weight which include low birth weight (Connor *et al.*, 2012), high birth weight (Lecoutre *et al.*,2016) or no differences in birth weight (Chen and Morris, 2009). The disparities in birth weight outcome can be attributed to differential impact of obese pregnancies on placental function, where obesity may increase placental growth to cause foetal overgrowth or reduce placental blood flow to limit foetal growth (Howell and Powell, 2017).

The incidence of a low birth weight due to the dietary manipulation employed in this study can potentially compromise adulthood metabolic phenotype. We therefore compared glucose homeostasis in male and female offspring of dams exposed to overnutrition or undernutrition. With differences in body weight no longer apparent in young adulthood (pnd 90), the animals were challenged with a glucose load to determine glucose homeostatic response in a timed fashion. The data on blood glucose AUC during the OGTT indicated that glucose regulation was well-controlled in these animals and no indication of glucose intolerance arising from the early-life nutritional insults. These findings suggest that low birth weight arising from maternal overnutrition or undernutrition prior to and during pregnancy may not be associated with glucose intolerance at 3 months of age in rats. In contrast, male mice from dams fed a high fat diet during pregnancy exhibited glucose intolerance at 2 months of age (Zhang *et al.*, 2019), while human studies on adults that were exposed to famine during the later stages of gestation showed higher glucose intolerance or type 2 diabetes than those born before or conceived after the famine (Roseboom *et al.*, 2001). This discrepancy may be due to differences in dietary components, window of exposure, specie and experimental design.

Unlike offspring of under-nourished dams who had normal glucose level under basal condition, the elevated baseline FBG level in over-nourished offspring may be an indication of a prediabetic state that could become apparent in the presence of a more severe metabolic stressor such as advancing age. Age-related decline in metabolic function has been reported in developmental programming studies. Progression from high insulin concentration at 3 months of age to development of full diabetes phenotype at 6 months has been reported in offspring of obese mothers (Samuelsson *et al.*, 2008). Similarly, a recent study showed age-dependent increase in most aspect of metabolic profile such as insulin, leptin, triglyceride and adiposity index in offspring of obese dams (Rodríguez-González *et al.*, 2019).

The development of hyperinsulinaemia in male rats exposed to maternal overnutrition or undernutrition, while sparing the female, may be due to sex hormone interaction on the developing foetus (Dunn *et al.*, 2011) and is an indication that maternal nutrition programmes changes to insulin level in a manner that is sex specific. Blood glucose response to insulin challenge was however divergent particularly in the male offspring, with development of insulin resistance in male rats exposed to maternal undernutrition, whereas offspring of over-nourished dams were sensitive to insulin. It is thought that low birth weight is a risk factor for insulin resistance (Cekmez *et al.*, 2013). While this was captured in the under-nourished group, offspring of over-nourished dams were not insulin resistant at 3 months of age. This suggests that birth weight index is a crude predictor of later life metabolic dysfunction. The induction of insulin resistance may be a corollary of hypertriglyceridaemia in male offspring exposed to maternal under-nutrition, which support of the close association between elevated triglyceride level and insulin resistance (Karpe *et al.*, 2011). Coupled with a weakened antilipolytic capability of insulin, accumulation of triglyceride-rich chylomicrons in insulin sensitive tissues alters insulin sensitivity (Karpe *et al.*, 2011). Similar to our results, others have also reported disturbance of lipid metabolism in rat offspring as early as third week of life (Zhu *et al.*, 2016).

A limitation to this study is a lack of data on the stress axis which can be activated by poor uterine environment to affect foetal outcome (Entringer *et al.*, 2012). Future studies will determine the mechanisms that underlie the differential metabolic phenotype reported in this study. Meanwhile, a recent study showed that metabolomic signatures of low birth weight are linked to insulin resistance and oxidative stress (Metrustry *et al.*, 2018).

In conclusion, we demonstrate that similar programming outcome on birth weight does not strictly translate to comparable metabolic phenotype in adulthood. Evidence of insulin resistance in male offspring of under-nourished dams supports our hypothesis that maternal undernutrition exerts a more debilitating metabolic effect in the offspring compared to maternal overnutrition.

## References

Akyol, A., Langley-Evans, S. C. and McMullen, S. (2009). Obesity induced by cafeteria feeding and pregnancy outcome in the rat. British Journal of Nutrition, 102(11): 1601–1610.

Barker, D. J. and Osmond, C. (1986). Infant mortality, childhood nutrition, and ischaemic heart disease in England and Wales. Lancet, 1: 1077–1081.

Barker, D. J., Winter, P. D., Osmond, C., Margetts, B. and Simmonds, S. J. (1989). Weight in infancy and death from ischaemic heart disease. Lancet, 2: 577–580.

Catalano, P. M., Presley, L., Minium, J. and Hauguel-de Mouzon, S. (2009). Fetuses of obese mothers develop insulin resistance in utero. Diabetes Care, 32(6): 1076–1080.

Cekmez, F., Canpolat, F. E., Pirgon, O., Aydemir, G., Tanju, I. A., Genc, F. A., Tunc, T., Aydinöz, S., Yildirim, S., Ipcioglu, O. M. and Sarici, S. U. (2013). Adiponectin and visfatin levels in extremely low birth weight infants; they are also at risk for insulin resistance. Eur Rev Med Pharmacol Sci. 17(4):501–6. PMID: 23467949.

Chaudhari, S., Otiv, M., Hoge, M., Pandit, A. and Sayyed, M. (2017). Components of Metabolic Syndrome at 22 years of Age - Findings From Pune Low Birth Weight Study. Indian Pediatr. 54(6):461–466. doi: 10.1007/s13312-017-1048-8. PMID: 28667716.

Dadi, A. F. and Desyibelew, H. D. (2019). Undernutrition and its associated factors among pregnant mothers in Gondar town, Northwest Ethiopia. PloS one, 14(4), e0215305. https://doi.org/10.1371/journal.pone.0215305.

Deputy, N. P., Dub, B. and Sharma, A. J. (2018). Prevalence and Trends in Prepregnancy Normal Weight - 48 States, New York City, and District of Columbia, 2011-2015. MMWR. Morbidity and mortality weekly report, 66(51-52), 1402–1407. https://doi.org/10.15585/mmwr.mm665152a3

Devaskar, S. U. and Chu, A. (2016). Intrauterine Growth Restriction: Hungry for an Answer. Physiology (Bethesda). 31(2):131–46. doi: 10.1152/physiol.00033.2015. PMID: 26889018; PMCID: PMC4895444.

Dunn, G. A., Morgan, C. P. and Bale, T. L. (2011). Sex-specificity in transgenerational epigenetic programming. Hormones and Behaviour, 59(3): 290–295.

Entringer, S., Buss, C., Swanson, J.M., Cooper, D.M., Wing, D.A., Waffarn, F. and Wadhwa, P.D. (2012). Fetal programming of body composition, obesity, and metabolic function: the role of intrauterine stress and stress biology. J Nutr Metabx; 2012:632548. doi: 10.1155/2012/632548.

Hanson, M. A. and Gluckman, P. D. (2014). Early developmental conditioning of later health and disease: physiology or pathophysiology? Physiology Reviews, 94(4): 1027–1076.

Howell, K. R. and Powell, T. L. (2017). Effects of maternal obesity on placental function and fetal development. Reproduction (Cambridge, England), 153(3), R97–R108. https://doi.org/10.1530/REP-16-0495.

Koletzko, B., Godfrey, K.M., Poston, L., Szajewska, H, van Goudoever, J.B., de Waard, M., Brands, B., Grivell, R.M., Deussen, A.R., Dodd, J.M., Patro-Golab, B. and Zalewski, B.M (2019). Nutrition During Pregnancy, Lactation and Early Childhood and its Implications for Maternal and Long-Term Child Health: The Early Nutrition Project Recommendations. Ann Nutr Metab 74:93–106. doi: 10.1159/000496471.

Labayen, I., Ruiz, J. R., Ortega, F. B., Loit, H. M., Harro, J., Veidebaum, T. and Sjostrom, M. (2010). Intergenerational cardiovascular disease risk factors involve both maternal and paternal BMI. Diabetes Care, 33(4): 894–900.

Metrustry, S. J., Karhunen, V., Edwards, M. H., Menni, C., Geisendorfer, T., Huber, A., Reichel, C., Dennison, E. M., Cooper, C., Spector, T., Jarvelin, M. R. and Valdes, A. M. (2018). Metabolomic signatures of low birthweight: Pathways to insulin resistance and oxidative stress. PloS one, 13(3), e0194316. https://doi.org/10.1371/journal.pone.0194316

NIH. DHHS, PHS; 1985. Guide for the Care and Use of Laboratory Animals. NIH Publication No. 85-23.

Odhiambo, J. F., Pankey, C. L., Ghnenis, A. B. and Ford, S. P. (2020). A Review of Maternal Nutrition during Pregnancy and Impact on the Offspring through Development: Evidence from Animal Models of Over- and Undernutrition. International journal of environmental research and public health, 17(18), 6926. https://doi.org/10.3390/ijerph17186926

Ramírez-López, M. T., Vázquez, M., Lomazzo, E., Hofmann, C., Blanco, R. N., Alén, F., Antón, M., Decara, J., Arco, R., Orio, L., Suárez, J., Lutz, B., Gómez de Heras, R., Bindila, L. and Rodríguez de Fonseca, F. (2017). A moderate diet restriction during pregnancy alters the levels of endocannabinoids and endocannabinoid-related lipids in the hypothalamus, hippocampus and olfactory bulb of rat offspring in a sex-specific manner. PloS one, 12(3), e0174307. https://doi.org/10.1371/journal.pone.0174307

Rodríguez-González, G. L., Reyes-Castro, L. A., Bautista, C. J., Beltrán, A. A., Ibáñez, C. A., Vega, C. C., Lomas-Soria, C., Castro-Rodríguez, D. C., Elías-López, A. L., Nathanielsz, P W. and Zambrano, E. (2019). Maternal obesity accelerates rat offspring metabolic ageing in a sex-dependent manner. J Physiol. 597(23):5549–5563. doi: 10.1113/JP278232. Epub 2019 Nov 11. PMID: 31591717.

Roseboom, T. J., van der Meulen, J. H., Ravelli, A. C., Osmond, C., Barker, D. J. and Bleker, O.P (2001). Effects of prenatal exposure to the Dutch famine on adult disease in later life: an overview. Mol. Cell. Endocrinol. 185, 93–98.

Samuelsson, A. M., Morris, A., Igosheva, N., Kirk, S. L., Pombo, J. M., Coen, C. W., Poston, L. and Taylor, P.D. (2010). Evidence for sympathetic origins of hypertension in juvenile offspring of obese rats. Hypertension 55: 76–82.

Segovia, S. A., Vickers, M. H., Gray, C. and Reynolds, C. M. (2014). Maternal obesity, inflammation, and developmental programming. Biomed Research International, 418975. DOI: 10.1155/2014/418975.

Tarry-Adkins, J. L. and Ozanne, S. E. (2017). Nutrition in early life and age-associated diseases. Ageing Researh Reviews, 39: 96–105.

Zhu, W. F., Zhu, J. F., Liang, L., Shen, Z. and Wang, Y. M. (2016). Maternal undernutrition leads to elevated hepatic triglycerides in male rat offspring due to increased expression of lipoprotein lipase. Mol Med Rep. 13(5):4487–93. doi: 10.3892/mmr.2016.5040. Epub 2016 Mar 23. PMID: 27035287.

Zhang, Q., Xiao, X., Zheng, J., Li, M., Yu, M., Ping, F., Wang, T., and Wang, X. (2019). A maternal high-fat diet induces DNA methylation changes that contribute to glucose Intolerance in Offspring. Frontiers in Endocrinology, 10, 871. https://doi.org/10.3389/fendo.2019.00871.

